# Bioengineered recombinant kisspeptins with extended half-life exhibit novel peripheral function in a large-animal model

**DOI:** 10.64898/2026.02.04.703727

**Authors:** Vijay Kumar Saxena, Prasanthi Medarametla, Ajit Singh Mahla, Raghvendar Singh

## Abstract

Kisspeptins are the small peptide products encoded by the KISS1 gene and physiologically exist in various isoforms of variable length. They are the central regulators of reproduction, being a prominent driver of GnRH hormone secretion. Additionally, they have emerged as an important peripheral therapeutic target for many metabolic diseases like diabetes, obesity, and polycystic ovary syndrome (PCOS). Despite their therapeutic potential, their utility is severely limited by their short half-life. We have rationally bioengineered two versions of native kisspeptins, which we named HSK-1 and HSK-2. HSK-1 (8kDa) and HSK-2 (13kDa) are derived from the fusion of the albumin-binding ZAG domain from *Streptococcus zooepidemicus* with KP-10 and KP-52 versions of kisspeptins (KPs), respectively. In vitro assays confirmed that the proteins were functionally active and triggered downstream signalling. Molecular dynamics simulations of the proteins revealed their structural features relative to the native kisspeptin isoforms. Both molecules demonstrated stable receptor engagement, and ligand-induced conformational changes were observed, suggesting receptor activation. HSK proteins demonstrated an extended half-life and mostly acted peripherally in young animals. They reduced peripheral luteinizing hormone levels in young animals, likely representing a previously unrecognized mode of peripheral kisspeptin action.

**Graphical Abstract:** 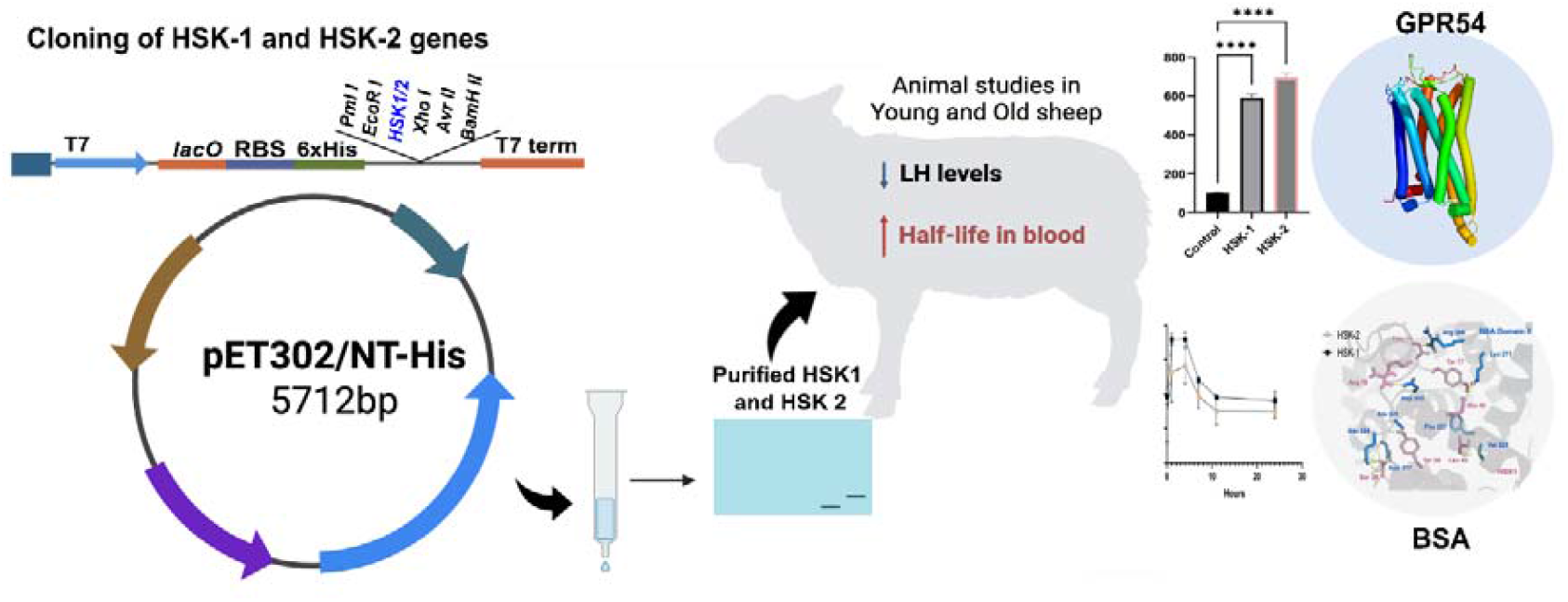

## 1.0 Introduction

Protein-based drugs have taken center stage, with a market share approaching 400 billion dollars annually, with hundreds of therapeutics either approved or in advanced clinical trials (Ebrahimi and Samanta, 2023). Recent advances in protein engineering have enabled the design of “smart,” stimulus-responsive biologics with enhanced functionalities and therapeutic potential.

Kisspeptin (KP), a small peptide hormone often hailed as a “magical molecule”, was first discovered for its role as a metastatic repressor of melanomas and hence named metastins (Lee et al., 1996). KPs are encoded by the *Kiss1* gene, which produces a 145-amino acid precursor protein that is cleaved into shorter active forms, including kisspeptin-54, kisspeptin-14, and kisspeptin-13 (Ohtaki et al., 2001). The ten-amino acid C-terminal structural motif (YNWNSFGLRF-NH2) is common among all kisspeptin isoforms and is considered essential for their activity.

KPs are produced by two major populations of neurons in the hypothalamus: the infundibular nucleus (analogous to the arcuate nucleus, ARC, in rodents) and the preoptic area (analogous to the rostral periventricular region of the third ventricle, RP3V in rats) (Rometo et al., 2007). These neurons project to activate hypothalamic gonadotropin-releasing hormone (GnRH) neurons, thereby stimulating GnRH secretion. KP receptors are G-protein-coupled receptors (GPR54), initially identified in the placenta (Ohtaki et al., 2001) and later localized in the brain, testis, ovary, liver, pancreas, and small intestine (Muir et al., 2001;Gutiérrez□Pascual et al., 2007). The binding of KPs to GPR54 triggers phospholipase C activation and the subsequent recruitment of secondary intracellular messengers, IP3 and DAG, which mediate intracellular calcium release. Additionally, DAG activates protein kinase C (PKC) and induces downstream phosphorylation of ERK1 and ERK2 signaling proteins.

KPs are key players in the onset of puberty and regulation of reproductive mechanisms, primarily by modulating GnRH (gonadotropin-releasing hormone) secretion from the hypothalamus. GnRH is a key hormone that drives the release of luteinizing hormone (LH) and follicle-stimulating hormone (FSH) from the pituitary gland (Seminara et al., 2003). They also participate in the negative feedback regulation of the hypothalamic-pituitary-gonadal (HPG) axis, mediated by sex steroid hormones such as estrogen and testosterone (Dorling et al., 2003). During the follicular phase of the cycle, pulsatile secretion of GnRH is regulated by the negative feedback effect of estradiol (E2). However, at the onset of the preovulatory stage, elevated E2 levels reverse and exert positive feedback, leading to a mid-cycle LH surge, inducing ovulation (Tsutsumi and Webster, 2009).

Emerging evidence indicates that KPs play an integral role in the local regulation of reproductive mechanisms across various peripheral reproductive organs, including the ovary, testis, uterus, and placenta (Cao et al., 2019). These findings emphasize their role in the peripheral reproductive process and mechanism, beyond the central hypothalamic controls. Marked KP alterations were observed in a mouse model of polycystic ovary syndrome (PCOS) characterized by very rapid, elevated in vivo LH pulsatility. (Esparza et al., 2020).

Beyond reproduction, emerging evidence indicates that KPs are involved in metabolic functions, linking energy status to reproductive processes (Manfredi-Lozano, Roa, and Tena-Sempere, 2018). There is significant evidence supporting the role of kisspeptin neurons in the arcuate nucleus (ARC) in transmitting metabolic signals to gonadotropin-releasing hormone (GnRH) neurons within the hypothalamic-pituitary-gonadal (HPG) axis (Bowe et al., 2012). KPs have been shown to enhance glucose-stimulated insulin secretion from both murine and human pancreatic islets without affecting basal insulin levels (Bowe et al., 2019). Moreover, the expression of KPs and their receptor, GPR54, is altered in the pancreas under metabolic stress conditions, such as high-fat diet (HFD) exposure and type II diabetes mellitus (T2DM). In addition, KPs have demonstrated beneficial and protective effects on liver metabolism, especially in relation to obesity (Song et al., 2014), T2DM (Dudek et al., 2016), and hepatic steatosis (Guzman et al., 2022).

Despite its therapeutic promise in reproductive and metabolic diseases, the clinical translation of KPs faces a major hurdle due to its extremely short biological half-life, i.e., approximately 4 minutes for KP-10 and 28 minutes for KP-54 in humans (Dhillo et al., 2005; Jayasena et al., 2011, 2015). The short half-life of the native KPs warrants frequent injections, which, in addition to logistical challenges, also risk receptor desensitization and compromise efficacy.

To overcome this challenge, we have engineered two recombinant kisspeptin fusion proteins that incorporate the ZAG domain from *Streptococcus zooepidemicus*. This was designed to extend the circulating half-life without compromising biological function. Here, we show that the recombinant KP fusion proteins remain functional both in vitro and in vivo and exhibit restricted penetration into the central nervous system in large-animal (sheep) models. This peripheral-selective approach constitutes a promising strategy to harness KP’s metabolic and reproductive effects without overstimulating the hypothalamic-pituitary axis.

## 2.0 Material and Methods

### 2.1 ethics Statement

Sheep breeds native to the Indian terrain (Malpura and Dumba) were used as animal models for the study. Sheep were stationed at an organized farm of ICAR-Central Sheep and Wool Research Institute, India, located in the semi-arid region of India (5°28 ‘E, 26°26 ‘N and altitude 320 m above sea level). All experimental procedures were approved by the Institute Animal Ethics Committee (IAEC) of ICAR-CSWRI, Avikanagar, Rajasthan, India.

### 2.2 Materials

LB broth base, IPTG, Fluo-3 AM, Ni-NTA agarose, and BL21 cells were purchased from Invitrogen (Thermo Fisher Scientific, USA). Bovine serum albumin, Urea, and Triton X-100 were procured from Sigma-Aldrich (Merck, USA). Anti-His-H8 antibodies (Thermo Fisher Scientific, USA) and anti-KP10 primary antibodies (Sigma-Aldrich, USA) were used in the study. Goat anti-mouse HRP cross-adsorbed secondary antibodies were procured from Southern Biotech (USA). Sheep LH ELISA kits were obtained from Abnova (Taiwan).

### 2.3 PCR-based fusion cassette generation and cloning

The fusion construct encoding the KP-52 region of the sheep Kiss-1 gene fused to the ZAG domain was commercially synthesized by GenScript (USA). The synthesized construct was cloned into the pcDNA3.1 expression vector. However, eukaryotic expression of the gene yielded unsatisfactory results; therefore, we opted to express the protein in a prokaryotic system. Accordingly, a PCR-mediated cassette was prepared to produce two protein versions: HSK-1 (KP-10 fused with the ZAG domain) and HSK-2 (KP-52 fused with the ZAG domain). The cassette was generated by PCR-aided amplification and fusion of the domain using custom-designed PCR primers, employing a megaprimer strategy. Primers AKpF1 and AKpR1 were used to amplify and prepare the HSK-2 cassette, whereas AKpF1 and BIJR2 were used for the HSK-1 cassette. The gene cassette, as depicted in **Fig. 1**, was cloned into the pET302 vector using directional cloning with EcoRI and XhoI restriction enzymes. The vector encodes an N-terminal His tag, and the cassette was cloned in-frame with this tag. All sequences and primers are provided as supplementary data. The vector was transformed into the DH5α strain of *E. coli*. Transformants were screened by colony PCR and restriction enzyme double digestion to confirm the presence of the insert. The cloned gene sequences were verified by sequencing and confirmed to match the designed cassette. Finally, glycerol stocks of the clones in DH5α *E. coli* were prepared for long-term preservation.

**Fig 1.**
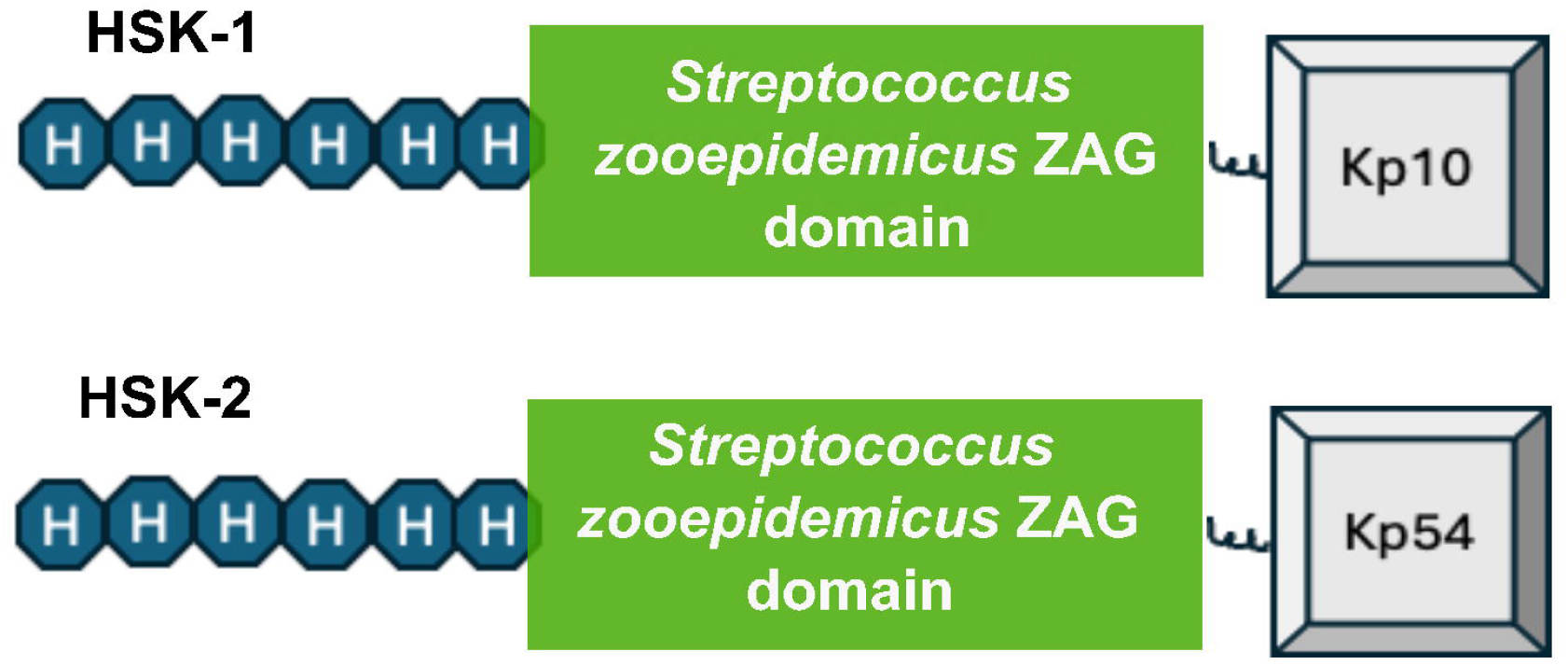
Schematic representation of the HSK fusion constructs. HSK-1 and HSK-2 each contain an Nterminal His tag fused to the Streptococcus zooepidemicus ZAG domain, followed by conjugation to different KP sequences, KP10 and KP52, respectively.

### 2.4 Expression and purification of recombinant kisspeptins

The recombinant proteins were expressed by transforming the prepared vector cassettes of HSK-1 and HSK-2 into the *E. coli* BL21 strain. The recombinant cells were bulk-cultured in Luria-Bertani (LB) medium containing ampicillin (100 µg/mL) at 37ºC and induced with isopropyl β-D-1thiogalactopyranoside (IPTG) at a final concentration of 1 mM for 4 hours. The culture was pelleted by centrifugation at 8,000 rpm for 20 minutes at 4ºC, and cell lysates were prepared in urea lysis buffer (8 M Urea, 20 mM sodium phosphate, 500 mM NaCl, pH 7.8) by gently rocking the cells at room temperature for 10 minutes. The cells were then sonicated on ice with brief pulses (5 pulses of 10 seconds each, with 20 seconds of rest between pulses). Cellular debris was pelleted by centrifugation at 3,000 rpm for 15 minutes, and the supernatant was collected for further purification.

Ni-NTA affinity chromatography was used to purify the recombinant proteins bearing an Nterminal hexa-Histidine tag. Ni-NTA resin (Thermo Fisher Scientific), having a high affinity and selectivity for 6×His-tagged proteins, was used according to the manufacturer’s denaturing purification protocol. The resin was equilibrated with 6 column volumes (CV) of binding buffer (8 M urea, 20 mM sodium phosphate, 500 mM NaCl, pH 7.8) and incubated with the lysate for 30 minutes at room temperature with gentle agitation. After binding, the resin was sequentially washed with 5 CV of binding buffer (pH 7.8), 5 CV of wash buffer (pH 6.0), and a final wash buffer (pH 5.3) to remove contaminants. The target protein was eluted using elution buffer (binding buffer composition, pH 4.0) and collected in 1 mL fractions. The purity of the eluted proteins was assessed by SDS-PAGE analysis.

### 2.5 Mild detergent-aided gradual dialysis of proteins

The proteins were purified as denatured proteins, then successfully refolded by slow dialysis under mild detergent conditions. 0.1% Triton X-100 was used as the detergent of choice to stabilize the proteins and maintain their solubility during gradual dialysis. The proteins were gradually dialysed in 50 mM Tris-HCl containing decreasing concentrations of urea (6, 5, 4, 3, 2, and 1 M Urea). Finally, the proteins were dialysed and stored in 50mM Tris-HCl containing 0.1% Triton X-100. The concentration of purified dialyzed concentrated protein was estimated by the Bradford method.

### 2.6 Electrophoretic Mobility Shift Assay

The peptide-protein binding assay was performed using native polyacrylamide gel electrophoresis (PAGE) under non-denaturing conditions. Bovine serum albumin (BSA) and recombinant proteins (HSK-1) were mixed in binding buffer (Tris-HCl, pH 7.5) at appropriate molar ratios and incubated at room temperature for 10 minutes to allow complex formation. Samples were loaded onto a 10% native PAGE gel and electrophoresed at 4°C in Tris-Glycine buffer. The gel was stained with Coomassie Brilliant Blue to visualize band shifts indicating peptide-protein complex formation. These shifts were analysed by comparing migration patterns with control samples. Peptide-only and protein-only reactions were included as controls to confirm specificity.

### 2.7 Modelling and Molecular Dynamic Simulations

#### a) HSK-1 and HSK-2 with GPR54 receptor

The GPR54 structure (PDB ID: 8XGS_A) was retrieved from the RCSB PDB database. The KP-10 peptide was extracted from the PDB structure (PDB ID: 8ZJD_F) and merged into the KP receptor structure (8XGS) following the alignment (this complex is referred as KP-10 from here onwards). HSK-1 and HSK-2 proteins were modelled in complex with the kisspeptin receptor (referred as HSK-1, HSK-2, respectively) using AlphaFold 3 server (Abramson et al., 2024). All four systems (Apo form, KP-10, HSK-1, HSK-2) were then prepared using Schrödinger’s protein preparation wizard. Here, hydrogens were added, and protonation states were generated at pH 7.4± 2.0 using PROPKA (Schrödinger, LLC, New York, NY, 2024-3). The structures were optimized and then minimized using the OPLS4 force-field (Lu et al., 2021). The systems were built to run the molecular dynamics simulations by adding the default POPC membrane (to helices). Systems were solvated using implicit solvent model TIP3P with the orthorhombic periodic boundary and buffer size of 15Å. OPLS4 force field was used to build the systems, and appropriate Na+/Cl− ions were added to neutralize the systems. Further, simulations were performed with Desmond GPU engine (Bowers, 2006). Simulations were run using the NPT ensemble at pressure 1.01325 bar and temperature 300K (using Martyna−Tobias−Klein barostat and Nosé−Hoover chain thermostat) with the default settings of RESPA integrator with timesteps 2, 2, and 6 fs applied for bonded, near, and far, respectively. The default Coulombic cutoff of 9 Å was applied and run for 1mS with 10 replicas using a random seed, resulting in a total of 10mS per system for HSK-1 and HSK-2. Apo form was run for 2.5mS (250ns*10 runs) and KP10 for 3.5mS (350ns*10 runs). Simulations were analysed using the Python scripts provided by the Desmond trajectory analyses from Schrödinger, unless otherwise stated (Supporting Information **S1-S3**). Protein-peptide interactions during the simulation were derived using the analyze_trajectory_ppi.py script. Further, the Desmond trajectories were converted to gromacs format using trj_no_virt.py script and Principal component analysis (PCA) was conducted using GROMACS trajectory analysis tools(Lindahl, Hess and van der Spoel, 2001). The results are visualized using PyMOL (The PyMOL Molecular Graphics System, Version 2.5.7 Schrödinger, LLC) with the help of the mode_vector.py script. All the simulation data was stored in Zenodo repository for the public use (**10.5281/zenodo.18221928**).

#### b) HSK-1 and HSK-2 with bovine serum albumin

The AlphaFold was used to predict the binding of HSK1 to bovine serum albumin. This preliminary structure was then used as a template to model the binding site of HSK1. The albumin chain (from the AlphaFold model) was replaced with the available crystal structure (PDB ID: 4F5S) following the structure preparation and alignment. The complex was simulated after the preliminary equilibration using the similar methods mentioned above and simulated for total of 4.5mS (500ns*9 replicates).

### 2.8 Intracellular Calcium Release Assay

Peripheral blood mononuclear cells (PBMCs) from sheep were used to test the effect of HSK-1 and HSK-2 on intracellular calcium levels. PBMCs were isolated from sheep blood using Histopaque according to the manufacturer’s protocol. A total of 10^5^ PBMCs were incubated with 20 µg of recombinant proteins, and 2 µM Fluo-3 AM was added in HBSS. Cells were incubated at 37°C for 20 minutes. An appropriate protein control was included in which Fluo-3 AM was added without protein, and an equivalent volume of the buffer in which the proteins were maintained following dialysis was included instead. After incubation, cells were washed three times with HBSS and then resuspended in 1 mL of HEPES-buffered saline. Steady-state fluorescence measurements were performed using a Tecan Infinite M200 spectrophotometer (Tecan Life Sciences, Switzerland). Fluorescence emission was measured at 526nm following excitation at 488nm. The reported fluorescence intensity represents the mean of four independent experiments on PBMCs.

### 2.9 Sandwich ELISA development for measurement of recombinant kisspeptin

Maxisorp Nunc plates (Thermo Fisher Scientific) were coated with 9□µg/mL of anti-human KP10 antibody (Sigma-Aldrich) in coating buffer at 37°C for 1 hour. The plates were then blocked with 3% bovine serum albumin (BSA) dissolved in TBST. Washing was performed three times using TBS containing 0.1% Tween-20 (TBST) with gentle shaking for 5 minutes each time. Fifty microliters (50□µL) of blood plasma samples and controls were added to the respective wells and incubated for 45 minutes. The washing step was repeated as described above. Anti-His H8 monoclonal antibody (Thermo Fisher Scientific) produced in mouse was added at a 1:3000 dilution and incubated for 1 hour. The plate was washed again three times with TBST. Goat anti-mouse IgG secondary antibody (cross-adsorbed against bovine, human, rabbit, goat, and rat IgG) was added at a 1:10,000 dilution and incubated for 1 hour at 37°C. The plates were washed, and 100□µL of ready-to-use TMB substrate was added for color development. The reaction was stopped with 0.16 M H□SO□, and absorbance was measured at 450 nm using a Tecan Infinite M200 spectrophotometer (Tecan Life Sciences, Switzerland).

### 2.10 Animal Experiments

#### a) Assessment of the biological half-life of the recombinant HSK-1 and HSK-2 in non-cyclic ewes

A total of six anoestrus adult Malpura sheep aged 2 to 6 years were taken for the study. Ewes in seasonal anestrus exhibit lower circulating levels of luteinizing hormone (LH), follicle-stimulating hormone (FSH), and estradiol, along with basal levels of progesterone, which reflect a suppressed reproductive state (Evans et al., 2001; Saxena et al., 2015), providing a clear baseline to assess the functional impact of recombinant kisspeptin. Two sheep received a subcutaneous injection of a 20-µg bolus dose of HSK-1, while the remaining four sheep received the same dose of HSK-2. Blood samples were collected prior to injection (0 h) and at 1 h, 4 h, 7 h, 11 h, and 24 h post-injection. Plasma was separated by centrifugation and used to measure recombinant kisspeptin and LH levels. Presence of recombinant kisspeptin in plasma was determined by sandwich ELISA as described earlier. The terminal phase was analyzed using a first-order, single-compartment decay model. A450 values from the terminal phase were natural log–transformed and plotted against time. Nonlinear regression analysis was performed using GraphPad Prism (version 10). The data were fitted to a monoexponential decay model assuming first-order kinetics:

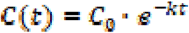

where C (t) is the measured absorbance at time t, Co is the estimated absorbance at the start of the terminal phase, and k is the apparent elimination rate constant. The apparent half-life (t_1/2_) was calculated by GraphPad Prism using the equation:

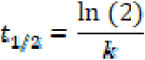

LH concentrations were determined using the Sheep LH ELISA Kit (Abnova) following the manufacturer’s protocol at 0 h and 1 h after protein administration.

#### b) Assessment of the LH level following HSK proteins administration in cycling adult ewes

Seven Dumba ewes (2-3 years of age) with normal estrous cycle were selected for the study. The regular cyclicity of these ewes confirmed functional HPG axis activity, making them suitable for evaluating LH responses following HSK protein administration. Six ewes received a bolus injection of HSK-1 at a dose of 20 µg, while one ewe was administered a 20 µg bolus dose of HSK-2The blood samples were collected at 0 h and 1h after the injection. The purpose of the experiment was to validate whether the HSK-1 and HSK-2 were able to cross the blood-brain barrier and modulate LH secretion as reflected by change in plasma LH levels.

### 2.11 Statistical Analysis

Except for data obtained from individual animals, all results are presented as mean ± standard deviation (SD). Statistical analyses were performed using Student’s *t*-test. A *p*-value < 0.05 was considered statistically significant.

## 3.0 Results

### 3.1 Production and characterization of recombinant DNA vector cassettes

Gene inserts coding for the fusion proteins were generated by the megaprimer PCR (Fig. **2a**), which yielded successful amplification of HSK-1 (225 bp) and HSK-2 (360 bp) gene constructs (Fig. **2b**). The genes were successfully cloned into the pET302 vector using the directional cloning strategy with EcoRI and XhoI as restriction enzymes. The transformants were screened by colony PCR and RE double digestion to release the insert (Fig. **2c**). The sequences were verified to be in the correct configuration and reading frame.

**Fig 2.**
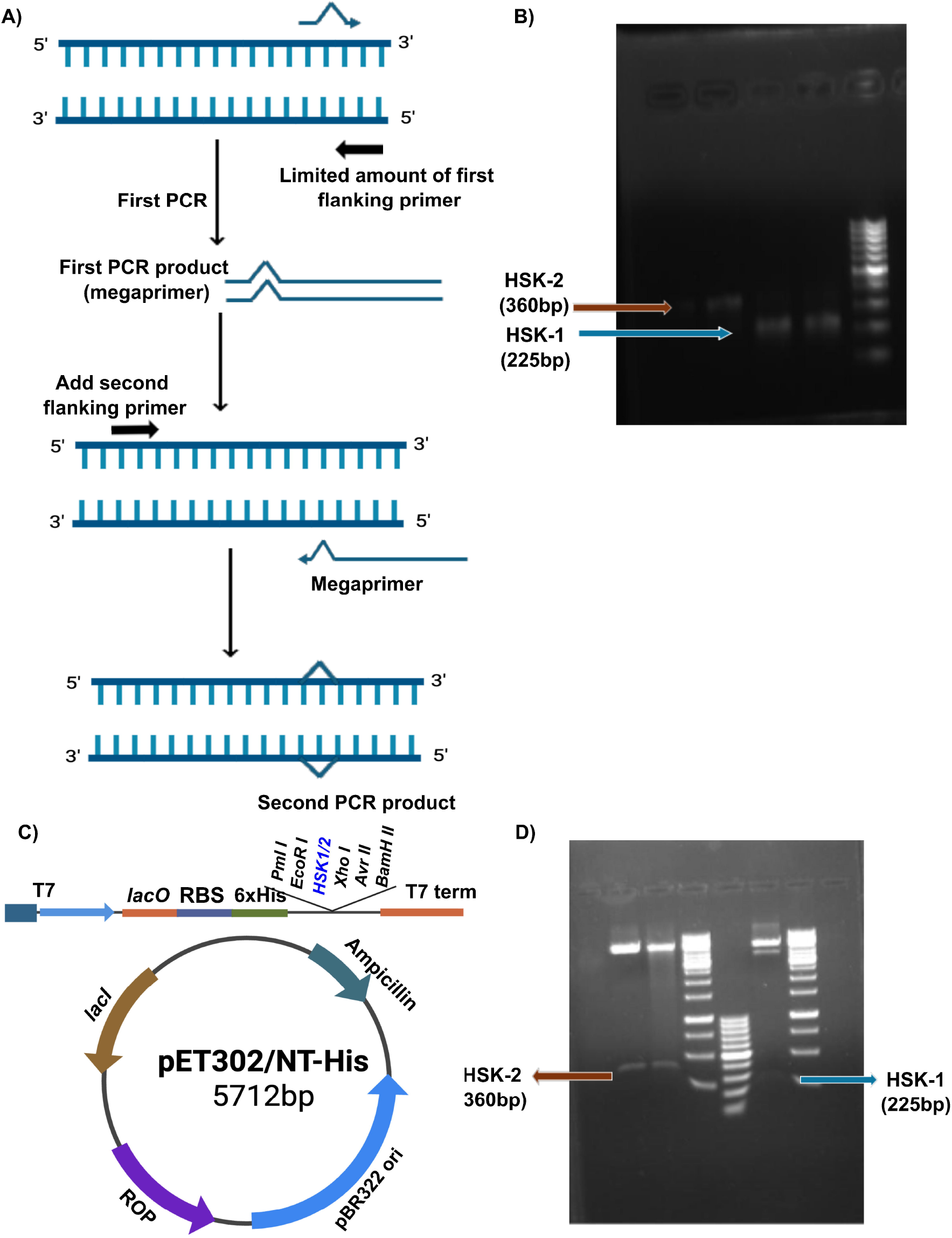
Cloning Strategy employed to construct cassette encoding fusion proteins. A) Schematic representation of megaprimer strategy B) PCR amplified product and their sizes of respective HSK1 and HSK-2 cassette C) Vector cassette map with cloned genes D) Restriction digestion-based confirmation of the release of the insert of HSK-1 and HSK-2

### 3.2 Production and purification of recombinant engineered KPs (HSK-1 and HSK-2)

Recombinant HSK-1 and HSK-2 proteins were successfully expressed in *E. coli* BL21 cells following IPTG induction. Both HSK protein versions were efficiently purified under denaturing conditions using Ni-NTA affinity chromatography and subsequently refolded via stepwise urea dialysis in the presence of 0.1% Triton X-100. The final dialyzed proteins remained soluble and structurally intact without any aggregation. The obtained yields were 3.2 mg/mL for HSK-1 and 2.8 mg/mL for HSK-2 from 200 mL cultures. SDS-PAGE analysis of the purified eluted fractions demonstrated that both HSK-1 and HSK-2 were highly pure after Ni-NTA affinity purification and refolding (Fig. **3a** and **3b**) and were suitable for downstream functional applications.

**Fig 3.**
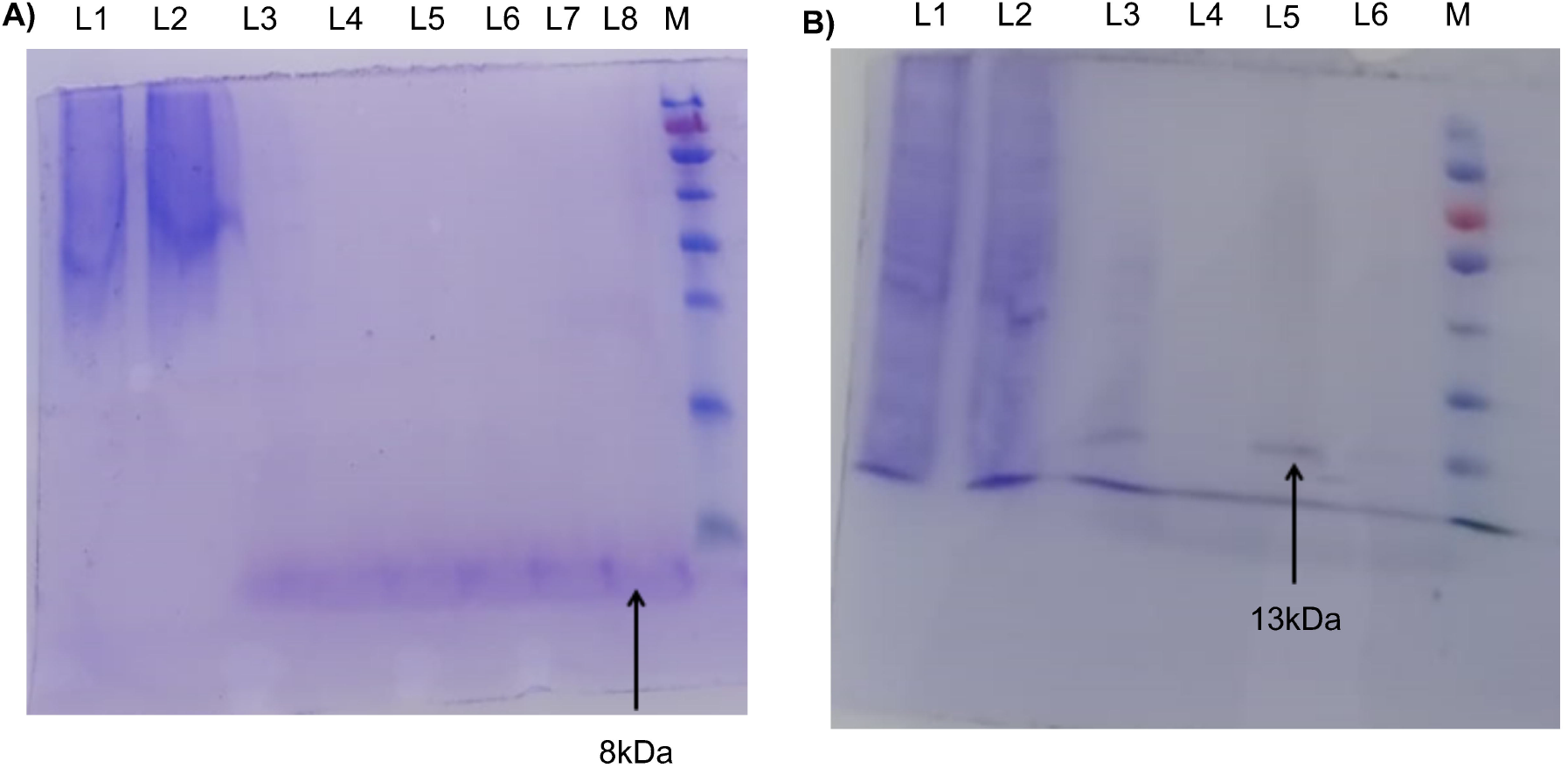
SDS PAGE analysis showing expression of full length HSK-1 (8kDa) and HSK-2 (13kDa). Lane information A) L1: Wash fraction, pH 6.0 L2: Wash fraction, pH 5.3 L3–L8: Purified HSK-1 fractions 1–6 Lane M: Molecular weight marker, with the lowest band at 10 kDa and the next band at 15 kDa Lane information B) L1: Wash fraction, pH 6.0 L2: Wash fraction, pH 5.3 L3–L8: Purified HSK-2 fractions 1–6 Lane M: Molecular weight marker, with the lowest band at 10 kDa and the next band at 15 kDa

### 3.3 HSK molecules bind BSA using the engineered ZAG domain

HSK-1 is expected to interact and bind to albumin due to the incorporated ZAG domain of *Streptococcus zooepidemicus*. To test this, the EMSA assay under native conditions was employed. A fixed amount of BSA (50 µg) was incubated with increasing concentrations of HSK-1. As shown in Fig. **4a**, the BSA band exhibited progressive retardation relative to the BSA-only control, indicating interaction and complex formation. HSK-1 alone, being a positively charged molecule, was not detected on the native PAGE gel, consistent with its electrophoretic mobility. Therefore, any observed shift in BSA mobility reflects interaction
with HSK-1. If no binding had occurred, the BSA band would have remained unchanged across all lanes.

**Fig 4.**
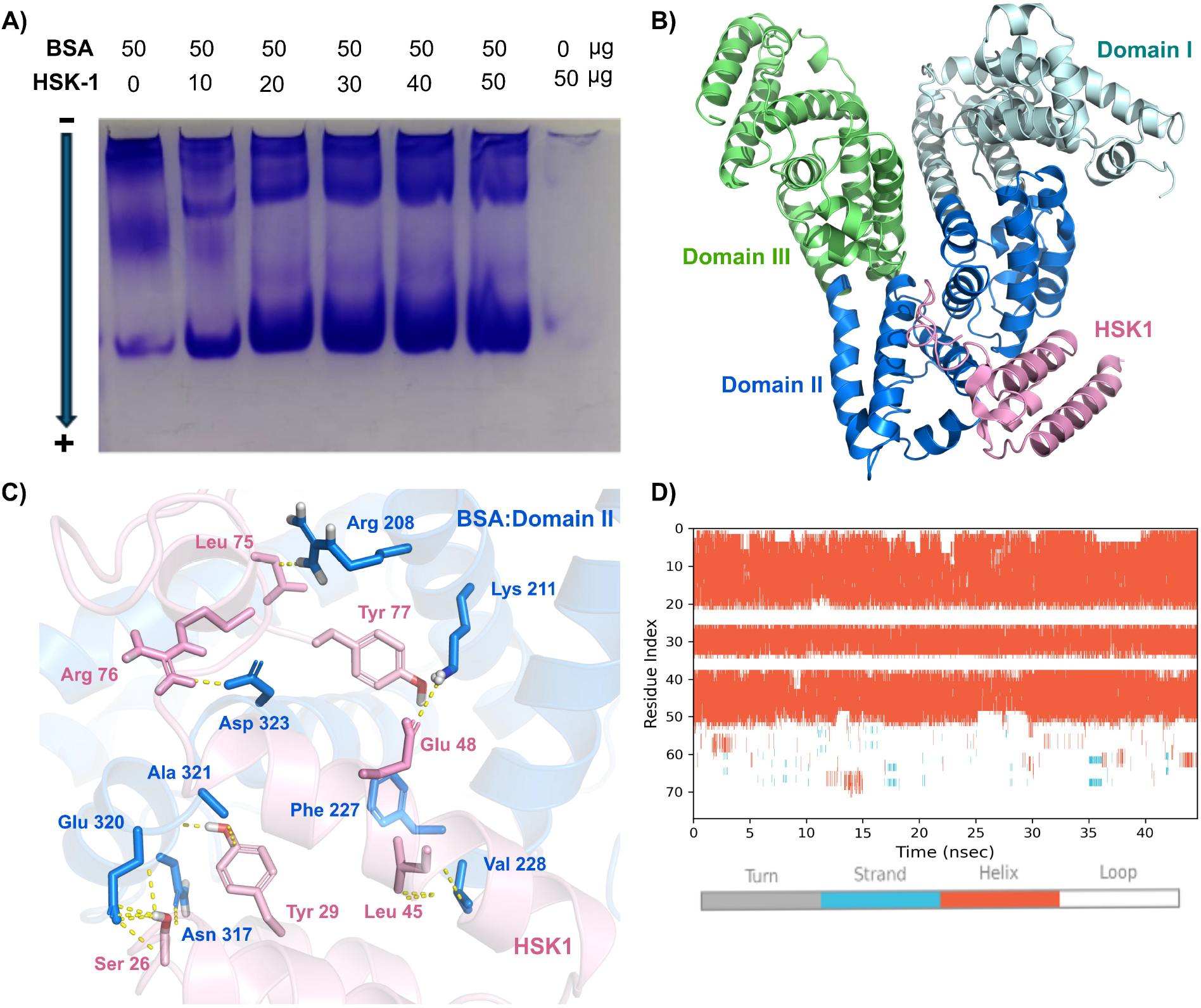
HSK molecules binds to albumins using their fused ZAG domains. A) EMSA showing interaction between HSK-1 and BSA. Increasing concentrations of HSK-1 cause progressive retardation of the BSA band on native PAGE, indicating complex formation. B) AlphaFold model of the Bovine Serum albumin (Domain I in Cyan, Domain II in Blue, and Domain III in Green) and the HSK1 peptide (pink); C) Interactions observed between the BSA Domain II (blue) and the HSK1 (pink)during the MD simulations, interactions are shown using yellow dashed lines; D) Composition of the secondary structural elements of the HSK1 peptide during the MD simulations

To further investigate this, AlphaFold was used to predict the binding of HSK-1 to bovine serum albumin. The AlphaFold prediction revealed that HSK-1 binds to the domain II (residues 196-383) of BSA, which is known as DrugSite1 (or Sudlow Site I; the binding site for several drugs such as warfarin, naproxen etc.) (Fig. **4b**). Interaction analysis revealed that the stability of the HSK-1 is maintained through hydrophobic interactions, hydrogen bonds, and salt bridges (Figure. **4c**). The Leu 45 and Tyr 29 residues of HSK-1 form hydrophobic interactions with the Val 228, Ala 321, and Phe 227 residues of albumin (domain II). These hydrophobic interactions are present for greater than 90% of the simulation time, indicating the ZAG domain interaction with BSA. The HSK-1’s Ser 26 and Tyr 29 formed hydrogen bonds with the Glu 320 (96% of simulation time) and Asn 317 (92% of time) respectively. Also, Glu 48 of HSK-1 formed salt bridges with the Lys 211, and Lys 37 with the Glu 226 for more than 50% of the time. These results further support the binding of HSK-1 to albumin through the Streptococcus zooepidemicus ZAG domain. During the simulations, the HSK-1 was stable in its secondary structure most of the time until residue 53; whereas the kisspeptin’s signature sequence (YNWNSFGLRF) adopted the loop conformation (Figure. **4d**).

### 3.4 HSK molecules functionally initiate downstream signalling

We hypothesized that the effects of HSK-1 and HSK-2 on intracellular Ca^2^□ levels would indicate whether these proteins can activate the intracellular signalling pathway of kisspeptin via their Cterminal decapeptide domain, which interacts with GPR54. Fluo-3AM is a membrane-permeable analog of Fluo-3 and is used as a calcium indicator. Fluo-3 AM ester itself does not bind Ca2+, but it is readily hydrolysed to Fluo-3 by endogenous esterases once the dye is inside the cells. There was a nearly sixfold increase in the fluorescence intensity of cells treated with HSK-1 and HSK-2 in comparison to untreated controls (Figure. **5a**), signifying the intracellular release of Ca^2^□within cells

### 3.5 Molecular Dynamics Simulations confirm the binding of HSK proteins to Kisspeptin receptors

The kisspeptin receptor apo form without KP-10 (Apo) was used as the reference structure to inspect the binding mode of the HSK-1 and HSK-2 proteins (Figure. **5b**). The protein transmembrane domains (TMs) were in the closed state in the apo form (Apo) compared with the other peptide-bound forms. The highest RMSF >5 Å was observed in the regions of the intracellular loop 3 (ICL3: connecting the TM5-TM6), TM6, and extracellular loop 3 (ECL3). Moderate fluctuations were observed in ECL2 (3-5Å). However, they are not as extensive as the ICL3, TM6, and ECL3 (Supporting Information Figure **S3**). Further, Principal component analysis (PCA) also highlighted large-scale movements of Intracellular loop 3 (ICL3). While TM5 remained stable in the Apo form and KP-10, binding of HSK-1 and HSK-2 induced conformational changes, where TM5 underwent kink transformation to accommodate the HSK’s C-terminal peptide region (Figure **5c**). This conformational change might be the reason for the strong binding of the HSK peptides to the KP receptor. Also, the HSK-1 and HSK-2 systems have undergone substantial rearrangements within the transmembrane domain to reorganize the C-terminal peptide region (Figure. **5c** and **S4**).

**Fig 5.**
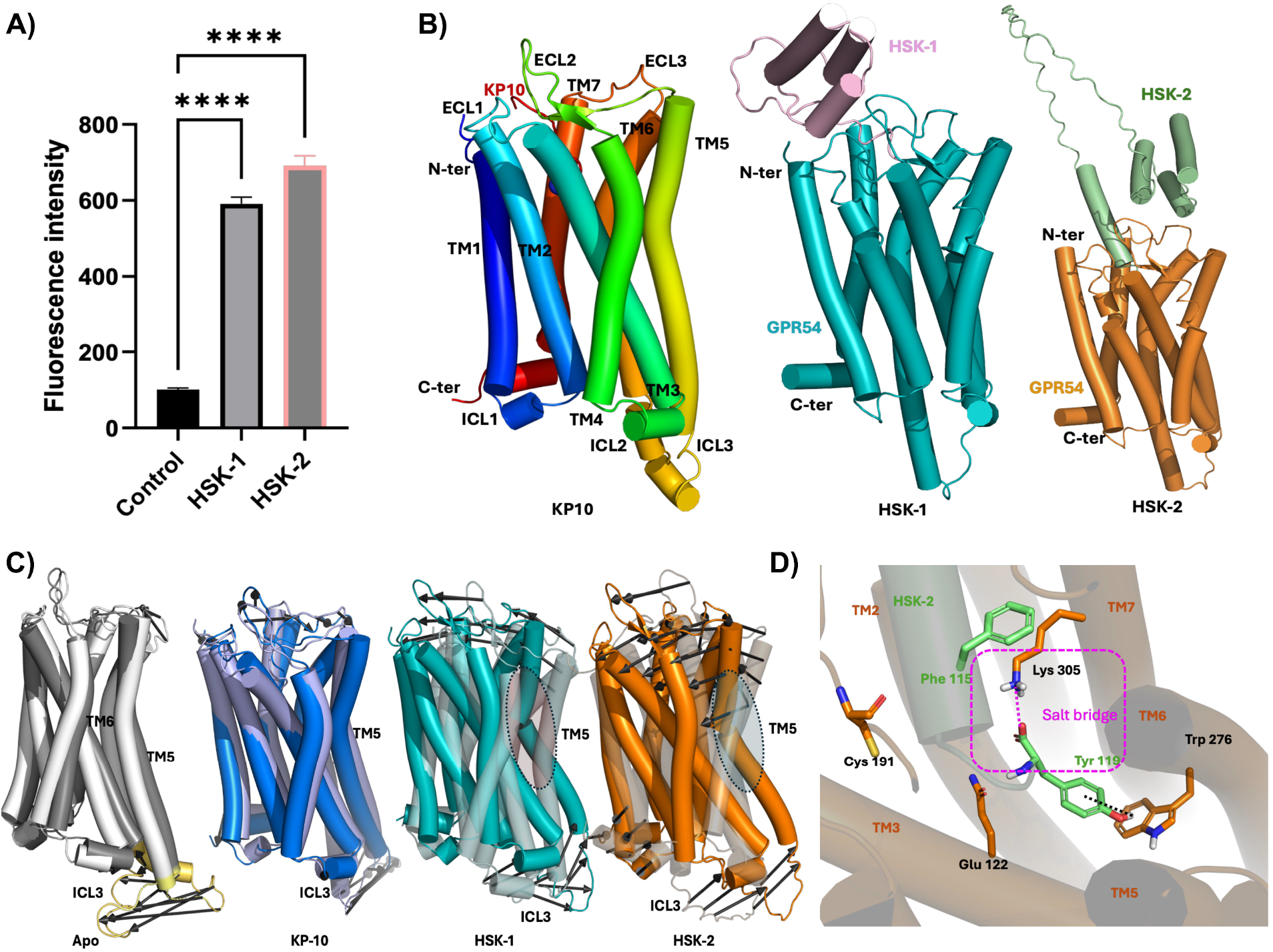
HSK molecules binds functionally with GPR54 to steer downstream signaling. A) Intracellular Ca^2^□ mobilization in response to HSK-1 and HSK-2. Cells loaded with Fluo-3 AM showed an approximately sixfold increase in fluorescence intensity following treatment with HSK-1 and HSK-2 compared to untreated controls, indicating activation of intracellular calcium signalling. B) Protein models used to investigate the binding of HSK peptides with the Kisspeptin receptor. Kisspeptin receptor (GPR54) with kisspeptide-10 (KP10: rainbow); HSK-1 (GPR54 with HSK-1 protein); HSK-2 (GPR54 with HSK-2 protein. C) Principal component analysis of the MD trajectories. Large conformational changes are indicated with dark arrows. TM5 and ICL3 which showed significant movements are highlighted using circles. D) The salt-bridge interactions (magenta lines) observed solely among the HSK-2 peptide’s Tyr 119 and Lys 305 of GPR54. Other residues involved in protein-peptide interactions are displayed as sticks (Phe 115-Lys 305: pication interaction; Tyr 119 and Trp276 hydrogen bond interaction; Glu 122 is also involved in salt bridge interactions).

Protein-peptide interactions were inspected further throughout the simulations. The functional peptide region YNWNSFGLRY has shown consistent interactions with the active site. Although several interactions, such as pi-cation, hydrogen bonds, and salt bridges, were observed, peptides are stabilized in the pocket mainly due to the hydrophobic interactions across all systems (KP10, HSK-1, and HSK-2). These hydrophobic interactions were observed with residues such as Ile 279, Val 118, Val 126, Tyr 302 and 190, Pro 106 and 296, Leu 202 and 283, and Glu 122 for greater than 80% of the time. Salt bridges were also observed with residues such as Lys 305, Glu 193, and Gln 201 (Figure 5d). Interestingly, Lys 305 has shown stable salt bridges in HSK-2 (Tyr 119 for 40% of the simulation time) compared with HSK-1 (1%). KP-10 didn’t show any salt bridges with Lys 305. The interaction with the Lysine might be playing an important role in the strong binding of the HSK-2 peptide in comparison with the others. Also, the protein is held tight together between the Lys 305 and Cys 191 (Supporting Information Figure S5). Distance analyses show that the separation between these residues remains stable at approximately 16Å throughout the simulation in the HSK-2 system, indicating a consistent spatial arrangement that may contribute to its stabilization (Figure **S5**). Overall, these results indicate that HSK peptides engage the KP receptor in a similar manner as the KP-10, where hydrophobic interactions are dominant in the binding. Additionally, HSK peptides displayed salt-bridge interactions with the Lys305 that enhance the binding stability and promote receptor activation and downstream Ca ^2^□signalling.

### 3.6 *In vivo* animal experiments in a large animal model (Sheep) demonstrating functionality and enhanced half-life of HSK molecules

#### *a) In vivo* stability and elimination profile

*In vivo* stability of the recombinant HSK-1 and HSK-2 was evaluated following a single 20 µg subcutaneous bolus injection in adult anoestrus sheep. Anoestrous ewes were particularly used in experiments as they do not have a high variation in reproductive hormones, as the HPG (Hypothalamo-pituitary-gonadal) axis is comparatively quiescent. HSK levels in plasma were quantified at multiple time points using an in-house, standardized ELISA specific for detecting the His-tagged kisspeptin proteins. Both recombinant kisspeptins were rapidly absorbed into the circulation, with a clearly detectable increase in plasma levels within 1 h of subcutaneous administration (Figure. **6a**). Following HSK-1 injection, the plasma concentrations of detected HSK-1 rose rapidly and peaked between 1 and 4 h, remained elevated above baseline for up to 7 h. Thereafter, levels gradually declined and approached pre-injection values by approximately 11 h. Analysis of the elimination phase indicated an estimated plasma half-life of approximately 1.5 h for HSK-1 (Figure. **6b**), demonstrating a significant improvement over native KP-10, which has a reported plasma half-life of less than 10 min. In contrast to HSK-1, HSK-2 displayed a more consistent and sustained pharmacokinetic profile. Plasma levels continued to increase up to 4 h post-injection, declined more slowly than HSK-1, and remained above baseline level well beyond 11 h (Figure. **6a**). HSK-2 demonstrated a longer estimated plasma half-life of approximately 3.5 h (Figure. **6b**) in close concordance with its pharmacokinetic profile. The enhanced stability of HSK-2 could be attributed to its larger molecular size and structural features. Together, these findings demonstrate that both recombinant kisspeptins exhibit markedly longer plasma half-lives than endogenous kisspeptins, with HSK-2 showing superior in vivo stability relative to HSK-1.

**Fig 6.**
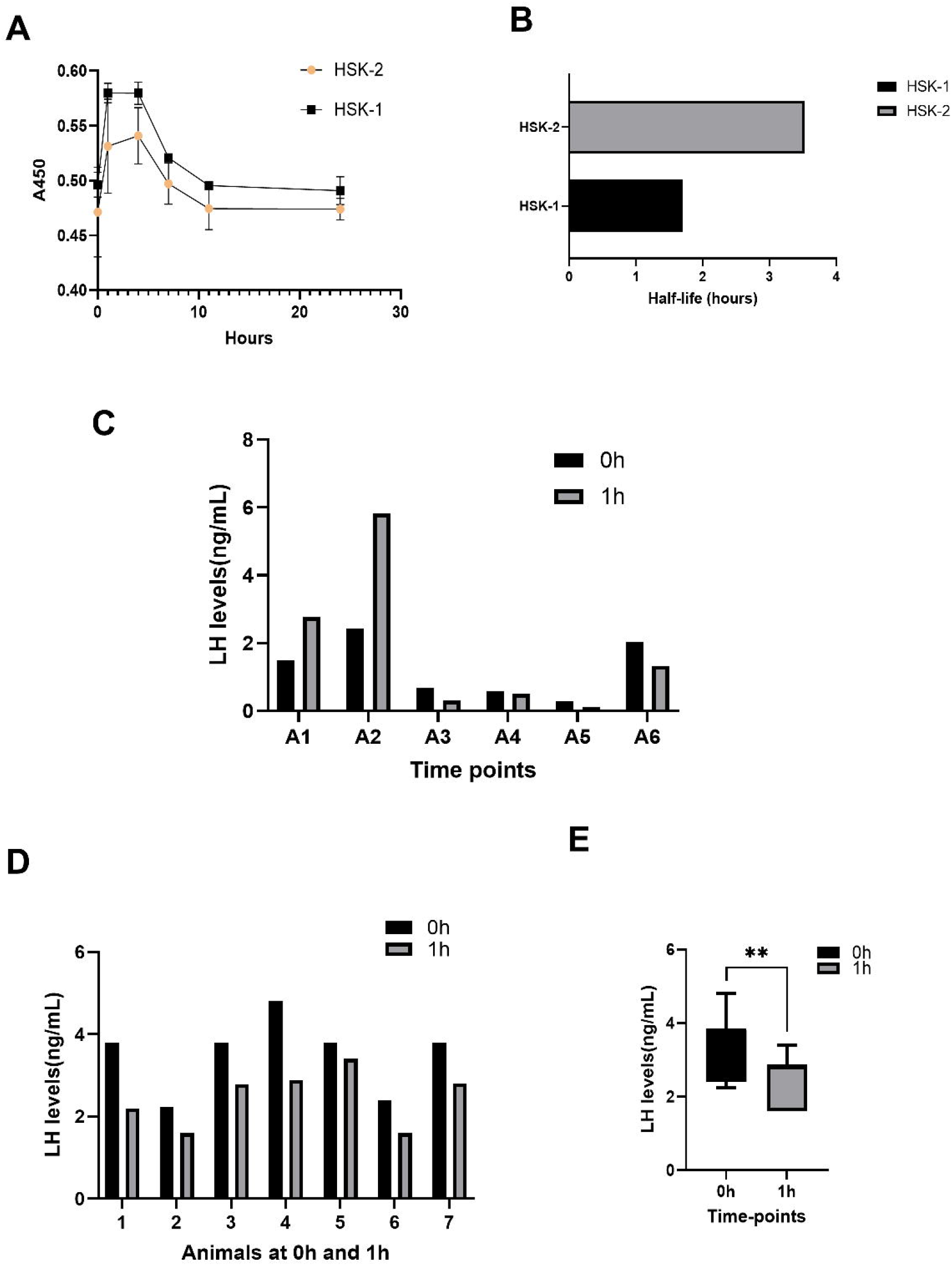
Animal experiments in sheep demonstrating increased plasma half-life and reduction in immediate peripheral LH levels on their treatment. A) Kisspeptin ELISA measuring recombinant kisspeptin absorbance in plasma and degradation profile after subcutaneous bolus injections B) Computed half-life of HSK-1 and HSK-2 proteins C) LH levels in plasma of Malpura anestrous sheep breed animals at 0h and 1h after the injections of HSK proteins. Animals (A1,A2,A3,A4) received bolus dose of HSK-2 and animals (A5,A6) received bolus dose of HSK-1. D) LH levels in plasma of experimental cycling young Dumba ewes at 0h and 1h after the injections of HSK proteins. Six young DUMBA sheep received a single bolus injection of HSK-1 (20 µg), while one animal received an equivalent dose of HSK-2 (20 µg). E) Mean LH levels were reduced significantly (p-value:0.002) at 0h and 1h after injection of HSK proteins in Dumba ewes.

#### b) Functional effects of recombinant HSKs on LH secretion from the pituitary

To assess the endocrine effects of recombinant kisspeptins, plasma luteinizing hormone (LH) concentrations were measured at baseline (0 h) and 1 h after administration of the proteins. A clear age-dependent divergence in LH responses was observed (Figure. **6c**). In older animals (A1 and A2; 6 years old), recombinant kisspeptin treatment increased circulating LH levels. Notably, one animal (A2) exhibited a robust response, with LH concentrations rising to more than two-fold above baseline. It signifies the stimulation of the gonadotrophs of the pituitary by the KP-GnRH system to release LH, which thereby increases the LH levels in plasma.

In contrast, all relatively younger animals (A3–A6) demonstrated a reduction in plasma LH concentrations following treatment. None of the younger sheep showed a surge in their LH levels or any evidence of gonadotropin stimulation. The absence of a positive LH response in these animals indicates that recombinant KPs failed to activate central GnRH signaling, which is likely due to their restricted permeability across an intact blood–brain barrier. Together, these findings demonstrate that age-dependent variability in response to recombinant kisspeptins, with central kisspeptin signalling associated with stimulation of gonadotropin release, is evident only in old animals.

#### c) Recombinant HSKs reduce plasma LH levels in Dumba sheep

To further investigate the unexpected suppression of plasma LH observed in young animals following recombinant kisspeptin treatment, the sample size was expanded. All animals included in this experiment were cycling and non-pregnant. Six young fat-tailed Dumba sheep received a single bolus injection of HSK-1 (20 µg), while one animal received an equivalent dose of HSK-2 (20 µg). Plasma samples were collected immediately prior to injection (0 h) and 1 h post-injection.

Across all young animals, administration of recombinant kisspeptins resulted in a significant reduction in mean plasma LH concentrations at 1 h compared with baseline levels (Fig **6d** and **6e**). This reduction in LH levels was consistently observed following administration of HSK-1 and HSK-2. Importantly, none of the treated young animals exhibited an increase in LH levels following administration of recombinant proteins.

## 4.0 Discussion

Kisspeptins (KPs) are biologically important neuropeptides derived from the proteolytic processing of the KISS1 gene precursor, yielding peptides of various lengths. Structural studies performed using NMR (Gutiérrez-Pascual *et al*., 2009) and circular dichroism spectropolarimetry (Saxena et al., 2011; Saxena et al., 2015; Thakuria et al., 2017) indicate that different kisspeptin fragments can adopt distinct conformations depending on their environment. They demonstrate environment-induced secondary structure changes that may influence receptor binding and activity. KPs are the central players in the physiological regulation of the onset of puberty, fertility and reproductive cycles in animals and humans (Trevisan et al., 2018; Hu et al., 2019; Sharma et al., 2020; Tsoutsouki et al., 2022; Xie et al., 2022) but have deeper peripheral roles which are being increasingly unravelled (Wolfe and Hussain, 2018; Izzi□Engbeaya and Dhillo, 2022; Tsoutsouki, Abbara and Dhillo, 2022; Patel et al., 2024; Sliwowska et al., 2024). KPs are modulators of pancreatic islet function and glucose homeostasis and are gaining recognition as key targets in diabetes management (Bowe et al., 2012; Schwetz, Reissaus, and Piston, 2014; McIntyre et al., 2019; Sridhar et al., 2025). They also demonstrate a protective effect on hepatic metabolism through their antioxidant properties (Hou et al., 2017). Moreover, stimulation of kisspeptin receptors has been demonstrated to attenuate liver steatosis and fibrosis in mouse models (Guzman et al., 2022).

All natural kisspeptin peptides (KP54, KP14, KP10) require a C-terminal RF-amide motif for receptor activation and functional activity. To counteract the short half-life of natural kisspeptins, various synthetic agonists such as FTM080 (Tomita et al., 2008), *KISS1*-305 (Matsui et al., 2012; Asami et al., 2013), TAK-448 (MVT-602) (Abbara et al., 2020), and TAK-683 (Asami et al., 2014) have been developed. All these lead agonists incorporate variable modifications, such as amino acid substitutions, N-terminal modifications, and conformational constraints to enhance in-vivo half-life. Similarly, the C6 analogue has an N-terminal lipophilic motif and additional chemical modifications that improved its in vivo stability and pharmacokinetics. Despite advances in half-life via chemical modifications, some peptides require multiple doses to maintain efficacy, as evidenced by the limited duration of elevated LH levels (Hu et al., 2022). In addition, the inherent chemical complexity of these peptides increases production costs, creates formulation challenges, and limits manufacturing scalability.

In the present study, we have developed two versions of KP molecules (HSK-1 and HSK-2) engineered as a fusion protein to the ZAG domain of *Streptococcus zooepidemicus* connected via a highly flexible linker. We demonstrate their production and purification using the most convenient *E. coli*-based recombinant protein production system, which is highly scalable and cost-effective. We analyzed the *in vitro and in vivo* functionality of the produced molecules and identified that they mostly act peripherally in young animals. These were found to have enhanced *in vivo* stability owing to reduced renal clearance, driven by their binding to serum albumins via the ZAG domain. Additionally, we found an unexpected function: peripheral administration of these molecules reduced plasma LH levels in sheep. It is recognized that these molecules could play an important role as protein-based KP mimetic therapeutics with enhanced half-life.

Initial attempts to express the proteins in mammalian HEK293 cells were unsuccessful, and thus, we transitioned to an E. coli-based expression system. The recombinant proteins were predominantly insoluble and were therefore purified using Ni-NTA affinity chromatography under denaturing conditions. Proteins were refolded using a mild detergent–assisted, stepwise dialysis protocol. The albumin-binding capability of the incorporated ZAG domain was subsequently evaluated using an electrophoretic mobility shift assay (EMSA). The observation of concentrationdependent retardation of the BSA protein in native PAGE gel confirms the interaction between HSK-1 and BSA. The interaction between HSK-1 and BSA observed in vitro was further corroborated by molecular dynamics simulations. Alphafold-based predictions indicated that HSK-1 binds stably to domain II of bovine serum albumin. Domain II in BSA corresponds to Sudlow Site I and is a canonical drug-binding region. It is the presence of strong hydrophobic interactions, hydrogen bonds and salt bridges that stabilise the binding. Many of these stabilizing interaction mechanisms remained intact throughout most of the simulation time, suggesting stable binding between HSK-1 and BSA. Cantante *et*.*al*, 2017 demonstrated albumin binding domain (ABD) of the ZAG protein from *S*.*zooepidemicus* binds mouse, rat and human albumins with a nanomolar affinity (Cantante et al., 2017). We observed that the ABD domain incorporated into HSK-1 binds bovine serum albumin, and our *in vivo* results of increased half-life suggest the ability to bind ovine serum albumin as well. Nevertheless, it’s not surprising as the principal ligand-binding sites of serum albumin, particularly the Sudlow site I within domain II, are highly conserved across Bovidae. Bovine and ovine serum albumins share greater than 90% sequence identity, with almost complete conservation of the hydrophobic and polar residues that define the ligand-binding pocket.

After analyzing the ZAG domain moiety of the fusion protein and confirming its intended functionality, it was important to assess the functional properties of the incorporated KP domains in the HSK-1 and HSK-2 molecules. GPR54 is highly distributed in the pancreas, placenta, pituitary gland and spinal cord. It is considerably abundant in the hypothalamus, limbic system, and basal ganglia, as well as in the spleen, peripheral blood leukocytes (PBLs), testis, and lymph nodes (Zhu et al., 2022). GPR54 is a Gq/11-coupled G protein-coupled receptor (GPCR), and activation by kisspeptin stimulates phosphatidylinositol 4, 5-biphosphate hydrolysis, Ca2+ mobilization, arachidonic acid release, and ERK1/2 MAPK phosphorylation. Thus, it is prominently involved in inositol triphosphate (IP3) induced release of Ca++ from endoplasmic reticulum stores. Given that kisspeptin receptor signaling cascade leads to IP□ and DAG-mediated mobilization of intracellular calcium, changes in cytosolic calcium levels represent a direct and functional readout of receptor activation. Accordingly, intracellular calcium measurements using the calcium indicator Fluo-3AM were performed to assess downstream signaling following kisspeptin stimulation. HSK molecules increased intracellular calcium in peripheral blood leukocytes, thereby exhibiting their functionality. Additionally, we analyzed the mechanism of HSK binding to GPR54 receptors using molecular dynamics simulations. The apo form of the kisspeptin receptor (Apo), without KP-10, was used as a reference to understand HSK-1 and HSK-2 binding. Compared to the closed Apo state, both proteins induced notable conformational changes in TM5, ICL3, and TM6, suggesting that these structural alterations are important for receptor activation. Hydrophobic interactions are strongly relevant in the interactions of both recombinant KPs, but HSK-2 forms a strong salt bridge with Lys305 and additional contacts with Cys191, which explains its higher binding affinity. The observations show that receptor flexibility and specific residue interactions determine ligand strength and specificity. Overall, this provides a good understanding of the interaction mechanism and stabilizing forces for HSK molecules binding to kisspeptin receptors.

Native kisspeptin peptides are characterized by extremely short plasma half-lives, with KP-10 being cleared within minutes in both rodents and humans, while KP-54 displays only marginally longer persistence (≤30 minutes). In contrast, both HSK-1 and HSK-2 demonstrated half-lives of 1.5h and 3.5h, respectively, after a single subcutaneous bolus. The prolonged systemic half-life of HSK-1 and HSK-2 likely results from their increased molecular size and their interaction with serum albumin. Both factors are well known to extend plasma residence time mainly by reducing the renal clearance and further decreasing susceptibility to proteolytic degradation (Dennis et al., 2002).

In aged sheep, administration of recombinant kisspeptins resulted in increased LH levels, consistent with the well-established role of kisspeptin as a potent upstream activator of GnRH neurons. This response could be easily explained by age-related compromise of blood–brain barrier integrity, which has been observed throughout mammalian species and permits enhanced penetration of larger peptides into the hypothalamus (Chen et al., 2009; Montagne et al., 2015). Such blood–brain barrier alterations with ageing have previously been implicated in altered neuroendocrine signaling and increased central accessibility of circulating molecules (Banks, 2009).

In sharp contrast, young adult sheep exhibited a significant reduction of circulating LH levels within 1h of administration of HSK-1 or HSK-2. This effect was further confirmed in Dumba sheep, where HSK proteins reduced LH levels even in cyclic ewes. This suggests that the reduction in LH levels was not an individual-animal or breed-specific phenomenon. Importantly, the absence of LH stimulation in young animals strongly suggests that recombinant kisspeptins do not cross an intact blood–brain barrier, as their administration fails to activate the hypothalamic–pituitary– gonadal axis. Consequently, the reduction in circulating LH observed in this investigation is an important and previously unrecognised peripheral action of recombinant kisspeptins. One plausible explanation is that KPs may enhance peripheral utilisation or clearance of LH, possibly by increasing receptor-mediated uptake in peripheral target tissues, thereby lowering its measurable plasma concentrations within 1h of injections. This raises an interesting and probable prospect that even native KPs may exert previously overlooked peripheral regulatory effects on LH dynamics independent of central GnRH stimulation. The overlooked functional role is not surprising, as none of the earlier versions of injected KPs were peripherally selective.

Collectively, we developed two functional bioengineered kisspeptin molecules with an enhanced half-life. Our research suggests that these molecules reduce circulating luteinizing hormone (LH) levels. Comprehensive pharmacokinetic analyses and mechanistic studies examining LH regulation by these molecules, alongside native kisspeptin forms, will be important directions for future research.

## Acknowledgement

Authors thank the Director, ICAR-CSWRI Avikanagar, for all facilities and support for the conduction of experiment. Dr Kesavan Manickam is acknowledged for his advice on devising halflife calculations. Special acknowledgement to my late colleague, Dr. Davendra Kumar, whose vision and encouragement have always been a source of inspiration. Dr Narayanan Krishnaswamy is acknowledged for fruitful discussion and scientific brainstorming of ideas. The support of Suryaprakash, Vishnu, S.S. Rajput and Ramraj as skilled technical staff is acknowledged. The authors acknowledge CSC-IT Center for Science, Finland for computational resources. Prasanthi Medarametla is supported by the Finnish Cultural Foundation. Protein production and purification, in vitro experiments and animal studies were conducted at ICAR–CSWRI, while molecular dynamics simulations and modeling were performed at the University of Eastern Finland. Both institutions are hereby gratefully acknowledged.

## CRediT authorship contribution statement

**Vijay Kumar Saxena:** Writing – review & editing, Writing – original draft, Methodology, Investigation, Formal Analysis Conceptualization. **Prasanthi Medarametla:** Methodology, Investigation, Writing – review & editing **Ajit Singh Mahla:** Methodology, Investigation, Writing – review & editing **Raghvendar Singh:** Methodology, Writing – review & editing, Resources

## Declaration of generative AI and AI-assisted technologies in the manuscript preparation process

Authors declare use of ChatGPT for language editing and correction. After using this tool/service, the author(s) reviewed and edited the content as needed and take(s) full responsibility for the content of the published article.

