## Supplementary Information for "Bioengineered recombinant kisspeptins with extended half-life exhibit novel peripheral function in a large-animal model"

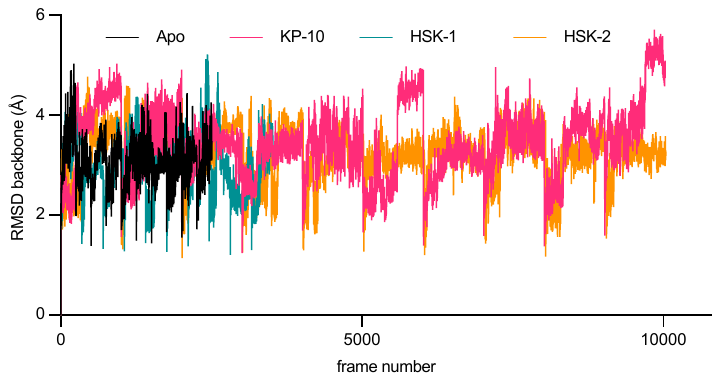

Figure S1. Route Mean Squere Deviations of the Kisspeptin in all the systems during the simulation. Apo (Kisspeptin:GPR54), KP-10 (GPR54 with Kisspeptide-10), HSK-1 (GPR54 with HSK-1), HSK-2 (GPR54 with HSK-2)

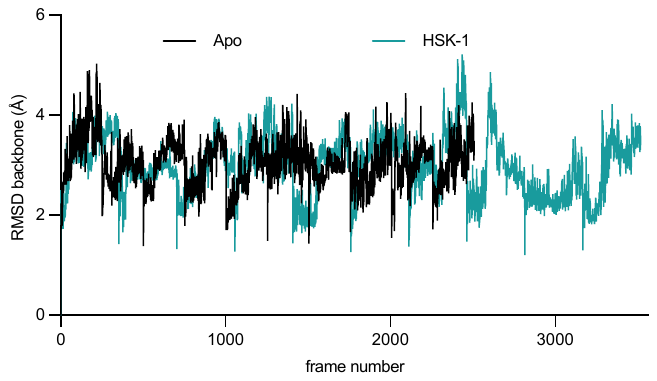

Figure S2. Route Mean Squere Deviations of the Kisspeptin in Apo system and HSK-1 from the figure S1 are depicted here for the clarity

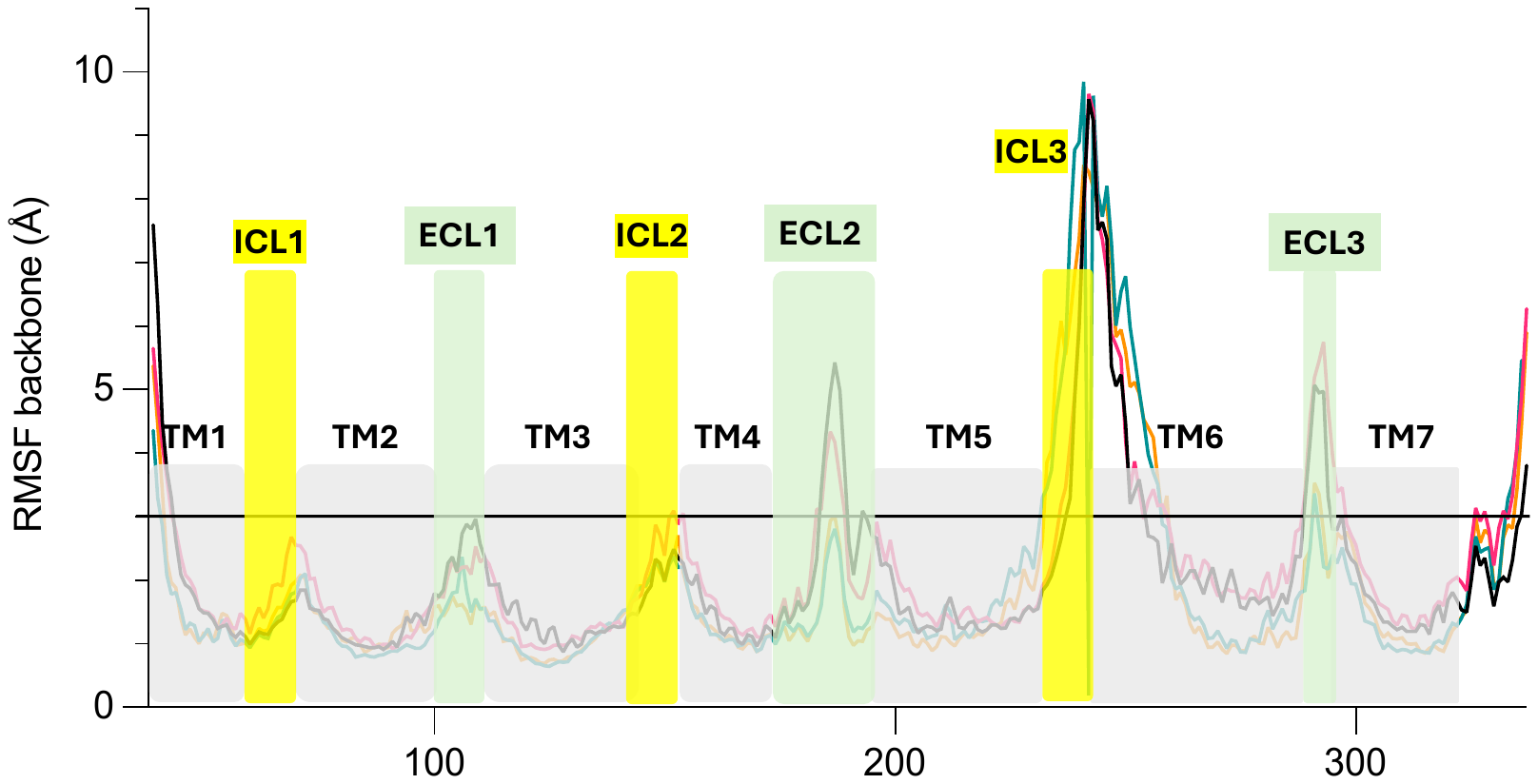

Figure S3. Route Mean Square Fluctuations (RMSF) of the systems during the simulation

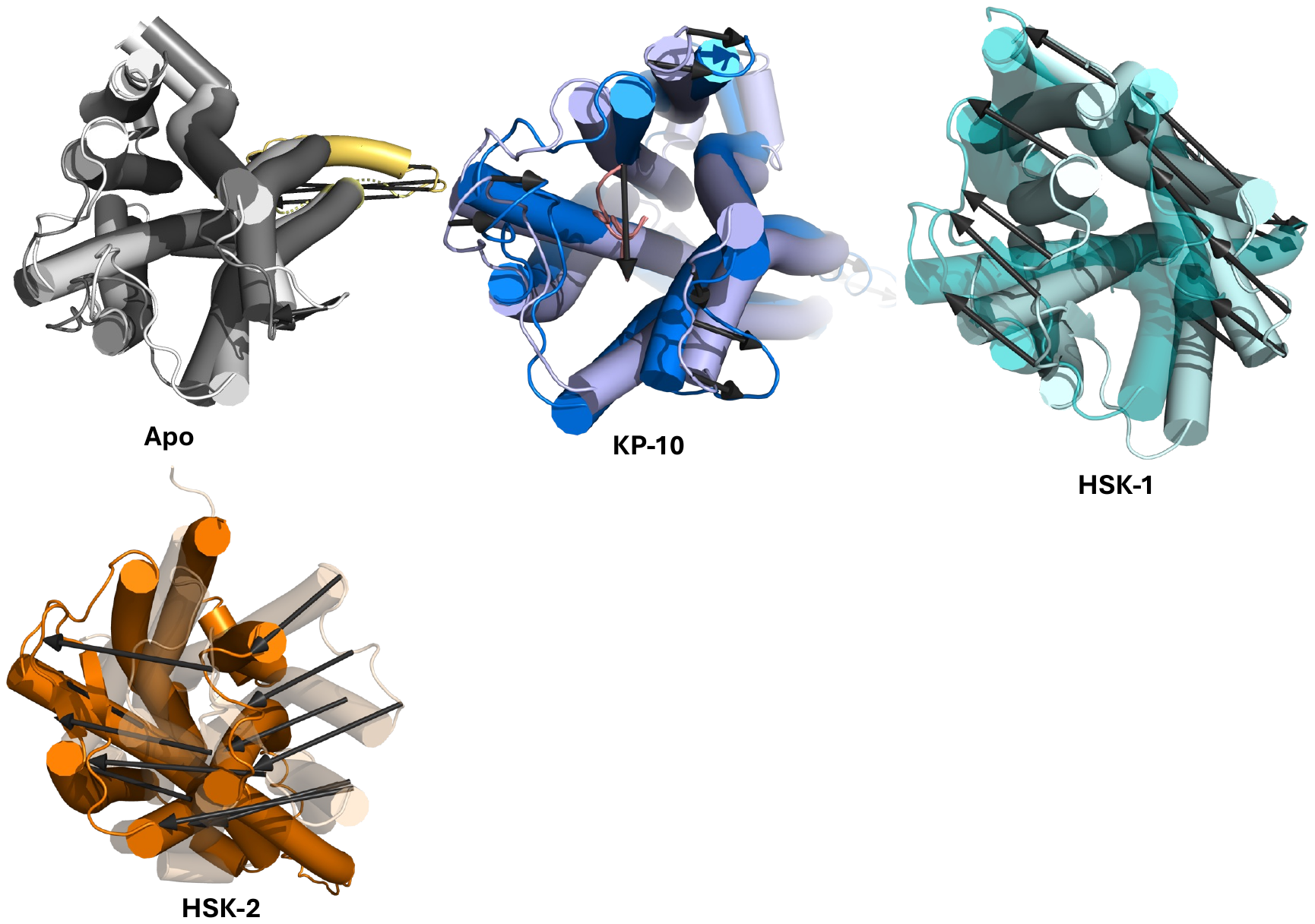

Figure S4. PCA analysis (top view) of the systems. Arrows indicate the movements during the MD

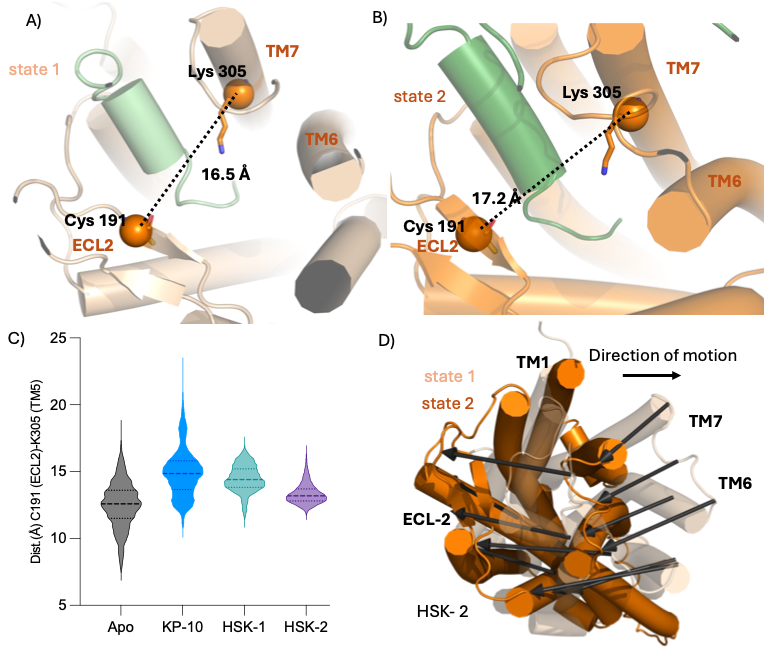

Figure S5. A and B) The distance between the residues binding the peptide in HSK-2 system of two states observed in the PCA analysis are shown. Ca atoms are shown as spheres. Kisspeptin receptor is shown as orange or light orange and HSK-2 peptide as green or forest green. C) Violin plot shows the distance between the Lys 305 and Cys 191 along the trajectory. The variation is not huge in the HSK-2 system providing the structural anchoring of these residues might be important for the stabilization of the complex. D) Reference picture of HSK-2 PCA analysis top view is shown here.
